# Bacterial community complexity in the phyllosphere penalises specialists over generalists

**DOI:** 10.1101/2023.11.08.566251

**Authors:** Rudolf O. Schlechter, Mitja N. P. Remus-Emsermann

**Affiliations:** Institute of Microbiology and Dahlem Centre of Plant Sciences, Department of Biology, Chemistry, Pharmacy, Freie Universität Berlin, Berlin, Germany; School of Biological Sciences and Biomolecular Interaction Centre and Bioprotection Research Core, University of Canterbury, Christchurch, 8011, New Zealand

**Keywords:** Phylloplane, Sphingomonas, Methylobacterium, Spatial pattern, Species interactions, Leaf surface

## Abstract

The leaf surface, i.e. the phylloplane, is an oligotrophic and heterogeneous environment due to its topography and uneven distribution of resources. Despite being a limiting environment, leaves host bacteria that are abundant and establish spatially-structured communities. However, factors that drive spatial distribution patterns are not well understood. Since leaf-associated bacteria can have beneficial effects to their host, understanding the rules of the community assembly can lead to novel strategies for crop protection. To investigate changes in population density and spatial distribution of bacteria in synthetic communities, we examined the behaviour of two prevalent bacterial groups in the *Arabidopsis thaliana* leaf microbiota: *Methylobacterium* spp. (specialists) and *Sphingomonas* spp. (generalists). We designed synthetic communities composed of two (S2) or three strains (S3) in a full factorial design and tested whether density and spatial structure of communities in S3 can be explained by pairwise comparisons in S2. Our results showed that specialists are more susceptible to changes in population densities and spatial distribution patterns than generalists, with lower densities and aggregation patterns when a specialist is in S3 than in S2. Additionally, pairwise comparisons were not sufficient to explain the observed patterns in S3, suggesting that higher-order interactions play a role in the resulting structure of complex communities at the micrometre scale.

## Introduction

A main challenge in understanding microbial communities is predicting the outcome of their assembly process. While bacteria typically follow relatively simple assembly rules in homogeneous environments ^1–3^, these rules fail to predict their behaviour in heterogeneous environments ^4^. In nature, heterogeneous environments are widespread and communities are the product of combinatorial effects that operate at the local scale ^5,6^. The leaf surface, or the phylloplane, is one such environment, and while bacteria are abundant on leaves, their distribution is impacted by topography, availability of resources, and microclimatic conditions ^7–9^. The distribution of bacteria on the phylloplane depends on their ability to populate restricted microenvironments ^10,11^, resulting in spatially-structured communities ^12,13^. Additionally, these bacterial communities are compositionally consistent at higher taxonomic ranks within plant species ^14^, indicating that predictable drivers are influencing the assembly process.

Generally, assembly of bacterial communities is influenced by neutral processes, environmental filtering, and species interactions ^15,16^. While neutral processes such as dispersal and ecological drift play a role in earlier stages of the assembly process ^17^, the non-random spatial organisation of natural bacterial communities on *Arabidopsis thaliana* suggests that deterministic factors such as leaf topography and species interactions also shape the assemblage of stable communities on the phylloplane ^12^. Previous studies have examined the effect of species interactions in the leaf environment, including antagonism and resource competition ^4,18–21^, as well as positive interactions ^22^. However, studies on the interactions between microbes at the micrometre resolution have been neglected. As a consequence, the inferred interactions are likely the emergence at spatial scales much larger than the scale that is relevant for said interactions, leading to averaging underlying patterns ^6,23^.

By contrast, the distribution of bacterial populations at fine-grained spatial resolutions better reflects their ecological roles on the phylloplane. Co-inoculation of fluorescently-tagged *Pantoea eucalypti* 299R (syn. *P. agglomerans* 299R, *Erwinia herbicola* 299R) and *Pseudomonas syringae* B728a onto bean leaves showed differential colonisation sites than those from near-isogenic mixtures ^24^. Additionally, mixed aggregates of these strains were highly segregated, where only 3% of cells were in direct contact with another, and death cells were commonly found at the interface between aggregates, suggesting that spatial organisations can also reveal species interactions at the micrometre scale. At a local scale, individual cells can interact and influence neighbouring cells at short distances ^25^. Co-aggregation between *P. eucalypti* and *P. syringae* in the bean phylloplane was observed at a 5–20 µm range; however, much less frequent than the aggregation between near-isogenic strains ^13^. At the community level, co-aggregation is common among different bacterial taxa and similarly ranges between 5–10 µm ^12^.

Pseudomonadota (syn. Proteobacteria) dominate leaf-associated bacterial communities of many plant species ^14,26–28^. Two particularly abundant co-existing bacterial groups on the phylloplane are the two alphaproteobacterial genera *Methylobacterium* and *Sphingomonas* ^29–31^. *Methylobacterium* (methylobacteria) are resource specialists that make use of methanol released as a byproduct of the plant cell wall metabolism ^32^. By contrast, *Sphingomonas* (sphingomonads) are resource generalists that utilise many different photosynthates leaching from leaves ^4,33–35^.

Methylobacteria and sphingomonads are consistently prevalent in leaf-associated microbial communities. However, the role of their metabolic differences in influencing microbe-microbe interactions in the phyllosphere has not been addressed yet. Understanding the consequences of species interactions is key to explain community dynamics and the assembly process. Species interactions are often challenging to infer from microbiome studies, as species co-occurrence can be a consequence of many mechanisms, such as mutualistic, neutral, or competitive interactions ^36,37^. Thus, a more detailed and controlled experimental approach is needed to investigate species interactions in the plant environment^38^. Therefore, to study the co-existence between and within *Methylobacterium* and *Sphingomonas*, we selected three methylobacteria and two sphingomonads for a detailed spatial analysis. Differentially fluorescent derivatives of those strains were co-inoculated onto otherwise axenic *A. thaliana* plants in two-or three-species synthetic communities (SynComs) in a full factorial design. The resulting population densities of the different strains were determined and their spatial distributions were investigated on leaves in combination with fluorescence microscopy and image cytometry.

## Materials and Methods

### Bacterial strains and media

The bacterial strains used in this study are listed in Table 1. *Methylobacterium* sp. Leaf85 (MeL85), *Methylobacterium* sp. Leaf92 (MeL92), and *Methylobacterium radiotolerans* 0-1 (Mr0-1) were grown in Reasoner’s 2A (R2A) media broth or agar at 30°C, while *Sphingomonas melonis* FR1 (SmFR1) and *Sphingomonas phyllosphaerae* FA2 (SpFA2) were routinely grown in Nutrient Broth (NB) or Nutrient Agar (NA) at 30°C. *E. coli* S17-1 was grown in lysogeny broth (LB) at either 30 or 37°C.

**Table 1.** Strains used in this study.

| Strain | Transposon | Growth condition | Reference |
| --- | --- | --- | --- |
| *Escherichia coli* S17-1 | n.a. | LB | 40 |
| *Sphingomonas melonis* Fr1 | n.a. | NA; R2A; MM + 0.5% Succinate | 41 |
|  | ::Tn5::mSc::GmR | NA; R2A; MM + 0.5% Succinate | 42 |
|  | ::Tn5::sYFP2::KmR | NA; R2A; MM + 0.5% Succinate | This study |
|  | ::Tn5::mTq2::TcR | NA; R2A; MM + 0.5% Succinate | 43 |
| *Sphingomonas phyllosphaerae* FA2 | n.a. | NA; R2A; MM + 0.5% Succinate | 44 |
|  | ::Tn5::mSc::GmR | NA; R2A; MM + 0.5% Succinate | 42 |
|  | ::Tn5::sYFP2::GmR | NA; R2A; MM + 0.5% Succinate | This study |
|  | ::Tn5::mTq2::TcR | NA; R2A; MM + 0.5% Succinate | This study |
| *Methylobacterium radiotolerans* 0-1 | n.a. | R2A; MM + 0.5% Methanol | 45 |
|  | ::Tn5::mSc::GmR | R2A; MM + 0.5% Methanol | This study |
|  | ::Tn5::mTq2::KmR | R2A; MM + 0.5% Methanol | This study |
|  | ::Tn5::mTq2::TcR | R2A; MM + 0.5% Methanol | This study |
| *Methylobacterium* sp. Leaf85 | n.a. | R2A; MM + 0.5% Methanol | 29 |
|  | ::Tn5::mSc::GmR | R2A; MM + 0.5% Methanol | This study |
|  | ::Tn5::sYFP2::KmR | R2A; MM + 0.5% Methanol | This study |
|  | ::Tn5::mTq2::KmR | R2A; MM + 0.5% Methanol | This study |
| *Methylobacterium* sp. Leaf92 | n.a. | R2A; MM + 0.5% Methanol | 29 |
|  | ::Tn5::mSc::GmR | R2A; MM + 0.5% Methanol | This study |
|  | ::105::sYFP2::KmR | R2A; MM + 0.5% Methanol | This study |
|  | ::Tn5::mTq2::KmR | R2A; MM + 0.5% Methanol | This study |

N.a.: Not applicable.

Minimal media (MM; 1.62 g/L NH_4_Cl, 0.2 g/L MgSO_4_, 1.59 g/L K_2_HPO_4_, 1.8 g/L NaH_2_PO_4_・2H_2_O, 15 g/L agar, with the following trace elements: 15 mg/L Na_2_EDTA_2_・H_2_O, 4.5 mg/L ZnSO4・7H_2_O, 3 mg/L CoCl_2_・6H_2_O, 0.6 mg/L MnCl_2_, 1 mg/L H_3_BO_3_, 3.0 mg/L CaCl_2_, 0.4 mg/L Na_2_MoO_4_・2H_2_O, 3 mg/L FeSO_4_・7H_2_O, and 0.3 mg/L CuSO_4_・5H_2_O ^39^) supplemented with either 0.5% w/v succinate or 0.5% v/v methanol was used in conjugation experiments depending on the recipient strain (Table 1) to counter select against the auxotrophic *E. coli* S17-1 donor strain. In addition, MM supplemented with different carbon sources was used to select for bacterial populations within SynComs (Table S1). Where appropriate, media were supplemented with antibiotics in the following concentrations: ampicillin (Ap), 100 µg/mL; gentamicin (Gm), 15 µg/mL; kanamycin (Km), 50 µg/mL; and tetracycline (Tc), 15 µg/mL.

### Conjugation

Bacterial strains tagged with constitutively-expressed fluorescent protein coding genes were developed by delivering a transposon that encodes for a single monomeric fluorescent protein gene and an antibiotic resistance cassette into their genome ^42,46^. Transposons were delivered through bi-parental mating of transposon-bearing plasmids as described previously^47^. Briefly, *E. coli* S17-1 equipped with pMRE plasmids were used as donor strains (Table S2). Donor strains were grown in LB at 37°C until early exponential phase, i.e., at an optical density at 600 nm (OD_600_) of 0.4–0.5, and recipient strains were grown in either R2A or NB at 30°C until stationary phase. Then, donor and recipient strains were harvested by centrifugation (2,000 × *g* for 5 min), washed in phosphate buffer saline (PBS, 0.2 g/L NaCl, 1.44 g/L Na_2_HPO_4_ and 0.24 g/L KH_2_PO_4_), and mixed in a 1:1 OD_600_ ratio. A final volume of 50 µL of the bacterial mix suspension was inoculated onto an LB agar plate supplemented with 100 mM MgCl_2_ and incubated overnight at 30°C. Finally, bacterial mixes were recovered, washed, resuspended in 1 mL PBS, and plated in selective media. Positive transconjugants were cleaned from the donor by at least three successive restreaks on appropriate selective media.

### Plant growth and inoculation

*Arabidopsis thaliana* ecotype Columbia (Col-0) seeds were surface-sterilised in a solution containing 50% v/v ethanol and 6% v/v H_2_O_2_ for 90 seconds, then thoroughly washed three times with sterile distilled water. Before sowing, seeds were stratified in sterile water at 4°C for at least 2 days.

To determine changes in population density and spatial distribution of bacteria on leaves, arabidopsis plants were grown in Magenta GA-7 tissue-culture boxes (Bioworld) containing sterile zeolite as an inert soil substitute, as described previously ^43^. Briefly, 90 g of finely ground zeolite was added to each box, autoclaved, and stored at room temperature until use. Before transferring seedlings into the tissue-culture boxes, 60 mL of sterile ½ strength Murashige-Skoog (MS; Duchefa) liquid medium (pH 5.9) was added to the boxes and let soak for 5 min. Stratified seeds were germinated on ½ MS agar media-filled tips. For this, pipette tips were filled with molten ½ MS (pH 5.9) 1% w/v agar and left to dry until the medium set. Aseptically, filled-tips were cut to 5 mm long pieces and placed upright into ½ MS 1% w/v agar media-containing plates. Individual seeds were sown on each tip, then plates were sealed, and placed into a plant growth chamber. Seven days after germination on tips, four seedlings were transferred into each tissue-culture box using flame-sterilised tweezers. Plants were grown in a Conviron A1000 plant growth chamber in short day conditions (11-h photoperiod, light intensity ∼120–150 µE s^-1^ m^-2^), 80% relative humidity, and at 22°C for 5–6 weeks or until plant developmental stage 1.11 ^48^.

For plant inoculation, bacterial strains were grown on plates for 2–5 days, depending on the strain, and a loop of bacteria was resuspended and washed twice in PBS. Bacterial suspensions were adjusted to an OD_600_ = 0.005 and mixed in equal ratio for each community in a full factorial design, resulting in three conditions: near-isogenic control (C), two-species community (S2), and three-species communities (S3) (Figure 1, Table 2). Then, each box was inoculated with 200 µL of bacterial suspensions with an autoclaved airbrush (KKmoon Airbrush Model T130A). Plants were sampled at 7 and 14 days post-inoculation (dpi) for colony-forming units (CFU) counts and microscopy (Figure 1). Bacterial density was determined by harvesting leaf material from inoculated plants into 1.7-mL microcentrifuge tubes using flame-sterilised tweezers and scalpel to remove root material. After plant fresh weight was determined, 1 mL PBS supplemented with 0.02% v/v Silwet L-77 was added into each tube. Samples were shaken in a bead mill homogenizer (Omni Bead Ruptor 24, Omni International) for two cycles of 5 min at a speed of 2.6 m s^-1^, and sonicated for 5 min (Easy 30 H, Elmasonic). CFU were determined by serial diluting the samples and plating on selective media for each population within a community (Table S1).

**Figure 1.**
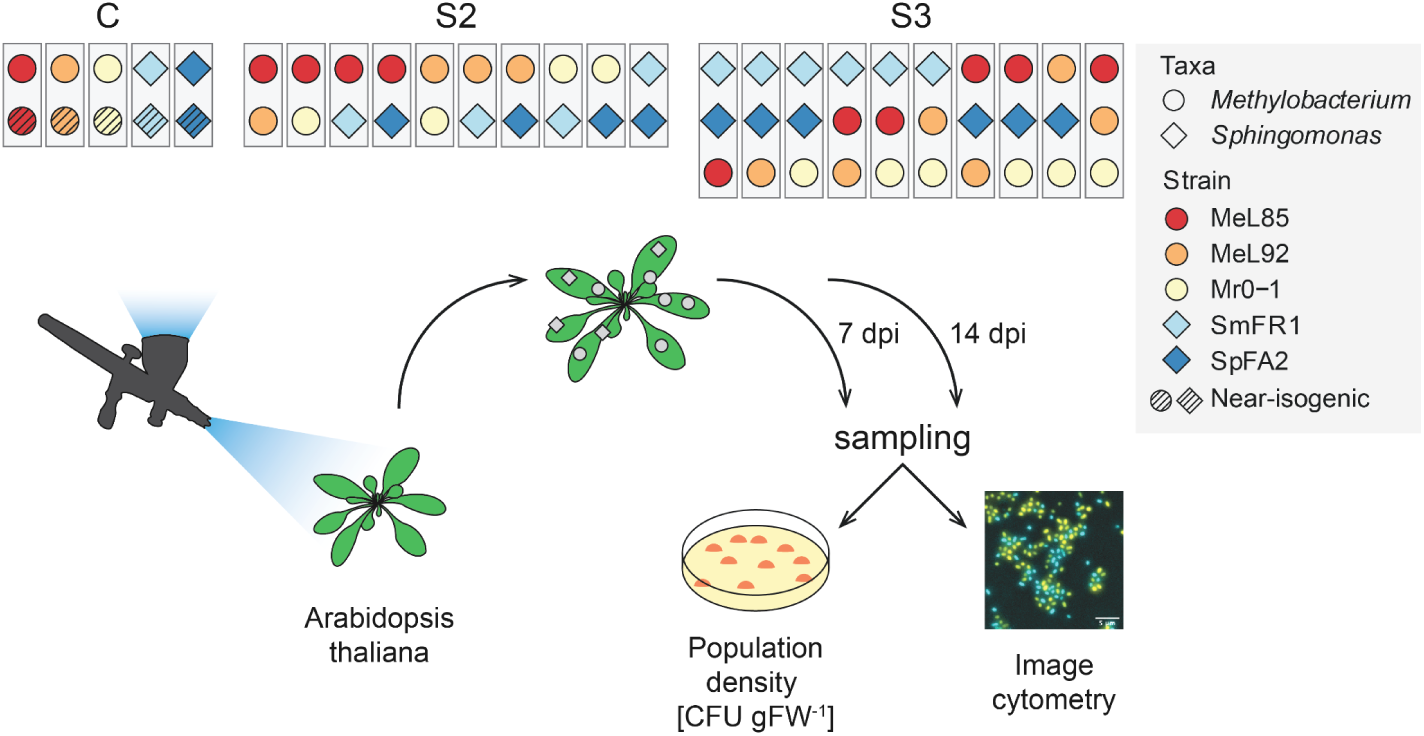
Experimental design. Four-week-old arabidopsis plants were airbrush-inoculated with a defined bacterial mix composed of either two near-isogenic strains (C), two different strains (two-species SynCom, S2), or three different strains (three-species SynCom, S3). Plants were sampled at 7 and 14 days post-inoculation (dpi) for CFU counts or image cytometry, as every strain expresses a unique fluorescent protein gene and selection marker. *Methylobacterium*: *Methylobacterium* sp. Leaf85 (MeL85), *Methylobacterium* sp. Leaf92 (MeL92), *Methylobacterium radiotolerans* 0-1 (Mr0-1). *Sphingomonas*: *Sphingomonas melonis* FR1 (SmFR1), *Sphingomonas phyllosphaerae* FA2 (SpFA2).

**Table 2.** Composition of Synthetic communities.

| Treatment | Community ID | Species 1 | Species 2 | Species 3 |
| --- | --- | --- | --- | --- |
| C | C.01 | MeL85 | MeL85 | n.a. |
|  | C.02 | MeL92 | MeL92 | n.a. |
|  | C.03 | Mr0-1 | Mr0-1 | n.a. |
|  | C.04 | SmFR1 | SmFR1 | n.a. |
|  | C.05 | SpFA2 | SpFA2 | n.a. |
| S2 | S2.01 | MeL85 | Mr0-1 | n.a. |
|  | S2.02 | MeL85 | MeL92 | n.a. |
|  | S2.03 | MeL92 | Mr0-1 | n.a. |
|  | S2.04 | MeL92 | SmFR1 | n.a. |
|  | S2.05 | MeL85 | SmFR1 | n.a. |
|  | S2.06 | Mr0-1 | SmFR1 | n.a. |
|  | S2.07 | SpFA2 | SmFR1 | n.a. |
|  | S2.08 | MeL85 | SpFA2 | n.a. |
|  | S2.09 | Mr0-1 | SpFA2 | n.a. |
|  | S2.10 | MeL92 | SpFA2 | n.a. |
| S3 | S3.01 | SmFR1 | SpFA2 | MeL85 |
|  | S3.02 | SmFR1 | SpFA2 | MeL92 |
|  | S3.03 | SmFR1 | SpFA2 | Mr0-1 |
|  | S3.04 | SmFR1 | MeL85 | MeL92 |
|  | S3.05 | SmFR1 | MeL85 | Mr0-1 |
|  | S3.06 | SmFR1 | MeL92 | Mr0-1 |
|  | S3.07 | MeL85 | SpFA2 | MeL92 |
|  | S3.08 | MeL85 | SpFA2 | Mr0-1 |
|  | S3.09 | MeL92 | SpFA2 | Mr0-1 |
|  | S3.10 | MeL85 | MeL92 | Mr0-1 |

N.a.: not applicable.

### Microscopy

To determine the distribution of SynComs on arabidopsis, leaf surfaces were sampled by cuticle tape lift as described previously ^12^. In brief, double-sided adhesive tape (1397 Orabond, Orafol) was attached onto microscope slides and the abaxial side of individual leaves of inoculated plants were placed onto the tape. Carefully, leaves were flattened and stripped off from the adhesive tape. Fixation was carried out by covering samples with 4% w/v formaldehyde solution (dissolved in PBS) for 2–3 h at room temperature. Then, the fixing solution was washed off by dipping the slides in PBS followed by sequential dehydration steps in 50%, 80%, and 100% v/v ethanol for 5 min each. Samples were dried using compressed air and stored at -20°C until processed.

Images were taken on a Zeiss AxioImager.M1 fluorescent widefield microscope at 1,000× magnification (EC Plan-Neofluar 100×/1.30 Ph3 Oil M27 objective) equipped with Zeiss filter sets 43HE, 46HE, and 47HE (BP 550/25-FT 570-BP 605/70, BP 500/25-FT 515-BP535/30, and BP 436/25-FT 455-BP 480/40, respectively), an Axiocam 506, and the software Zeiss Zen 2.3. Three-to-four independent plants, with at least two leaf samples per plant were used as biological replicates for each bacterial community. A minimum of 20 fields of view were acquired per leaf sample (124.94 µm × 100.24 µm) in up to three different fluorescence channels: red (43HE filterset), yellow, and/or cyan (46HE and/or 47HE filterset, depending on the SynCom, respectively, see Table 1); and phase contrast. Additionally, each field of view contained at least three images at different focus planes to increase the depth of field (0.5 µm step size) and to account for spherical aberrations.

### Image Analysis

Due to low signal-to-noise ratio, images were first processed in ImageJ/FIJI ^49^. Each stacked image was background subtracted (50 pixel, rolling ball) before applying a maximum intensity Z-projection. The contrast of the resulting image was enhanced (0.05%), and an unsharp mask filter was applied (5 µm radius, 0.5 stack). Maximum intensity Z-projections of each fluorescent channel were exported as individual images.

Bacterial cell segmentation was performed with omnipose using individual corrected fluorescent images as input and the “omni_bact” model for bacterial cell segmentation ^50^. After segmentation, labelled mask images were used to create binary pictures. Objects above 5 µm^2^ were filtered out in ImageJ/FIJI to exclude artefacts such as plant debris. Also, overlapping cells and bleed-through signals were corrected by subtracting the first channel (red) from the second channel (yellow or cyan). In the case of three-species SynComs, the resulting image was used to subtract from the third channel (cyan). Coordinates for each cell was determined as the centre of mass of each cell (Figure S1A). Population cell density was estimated as the sum of cells detected in each field of view per biological replicate (individual plant). The total area was estimated as the sum of areas according to the number of fields of view taken per biological replicate.

### Spatial Analysis

Spatial point pattern analysis was performed in R using the package *spatstat* ^51^. Cell coordinates in each field of view were converted into a point pattern and then combined into a hyperframe. Complete spatial randomness was evaluated with a quadrat-count χ^2^ test and a spatial Kolmogorov-Smirnov test on 500 random point patterns. In both cases, the null hypothesis is that the data is a realisation of a homogeneous Poisson point process.

The spatial association of bacterial cells within a population was estimated with the Ripley’s K-function for an inhomogeneous Poisson process at a given distance *r*, 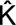(r). The estimate of 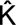(r) for each bacterial population was compared to a null model, which is the estimator K_inhom_(r) for an inhomogeneous Poisson point process using Monte Carlo simulation envelopes (n = 100) ^13^. Three spatial patterns can be detected at a given distance between points: aggregation (or clustering), regularity (or segregation), and randomness. Deviations between the empirical and theoretical K estimates suggest spatial aggregation (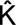(r) > K_inhom_(r)) or regularity (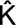(r) < K_inhom_(r)) within a population (Figure S1B). Spatial patterns of bacterial population in two-(S2) and three-species SynCom (S3) were compared to the spatial pattern of the respective near-isogenic populations (C).

The spatial association between two populations within a SynCom was estimated using the Ripley’s isotropic pair cross-correlation function for an inhomogeneous Poisson process at a given distance *r*, ĝ(r). Only point patterns that contained at least ten cells of each population to be analysed were considered. The estimate of ĝ(r) for each bacterial population was compared to the null model, which is the estimation of g_inhom_(r) for an inhomogeneous Poisson point process using Monte Carlo simulation envelopes (n = 100). Similarly to the K-function, three spatial patterns can be detected at a given distance between points based on pair correlations: co-aggregation (or clustering), regularity (or segregation), and randomness. Deviations between the empirical and null model suggest spatial co-aggregation (ĝ(r) > g_inhom_(r)) or regularity (ĝ(r) < g_inhom_(r)) between populations (Figure S1B). Spatial patterns between species pairs in S3 were compared to the spatial pattern of the respective pair in S2.

## Data analysis

Data analysis and graphical representations were performed in R. Homogeneity of variance was evaluated with the Breusch-Pagan test (*car* package). Normal distribution was evaluated with the Shapiro-Wilk test (*stats* package).

Population densities (either CFU gFW^-1^ or cell cm^-2^) were log_10_-transformed to compare between groups using a non-parametric two-samples Wilcoxon signed-rank test or a Kruskal-Wallis test, followed by a post hoc Dunn’s test for pairwise-comparisons. *P*-values were adjusted for multiple testing using the Holm method. Statistical tests were carried out using the package *rstatix*. The log_2_ fold change (log_2_FC) was calculated using the median of non-transformed data.

Pearson’s correlation was determined for the log_10_ transformed data of CFU gFW^-1^ and cell cm^-2^ with the *stats* package.

To analyse the results of 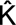(r) and ĝ(r) with replication, we calculated the frequency of a spatial pattern (aggregated, random, regular) by counting the occurrence of a spatial pattern at a given distance (Figure S1B) ^13^. From these plots, the absolute area under the curve (AUC) was calculated with the package *DescTools*, to obtain a magnitude of each spatial pattern. Then, fractional changes were calculated to evaluate the percentage differences of a spatial pattern for a strain or a strain pair in relation to a control (C for 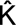(r) and S2 for ĝ(r), respectively). These data were analysed using either a one-sample Wilcoxon test (μ = 0 as reference), or with the non-parametric tests described above.

## Data availability

Images and raw data are available through Zenodo ^52^. Codes and analyses are available in the Github repository: https://github.com/relab-fuberlin/schlechter_phyllosphere_spatial_distribution.

## Results

### Changes in taxon-specific population density correlate with community complexity

We explored the impact of community complexity on bacterial population densities *in planta*, focusing on two prominent bacterial taxa, *Methylobacterium* and *Sphingomonas*. Employing a full factorial design (Figure 1), we evaluated bacterial population densities across three levels of community complexities: near-isogenic control (C), a two-species SynCom (S2), and a three-species SynCom (S3) at 7- and 14-days post-inoculation (dpi).

We first tested the influence of community complexity on individual bacterial populations at the CFU level, showing a pronounced and statistically significant effect (Kruskal-Wallis, *H*(2) = 307.02, *p* < 0.05). This effect was reflected by a significant 2.3-fold increase in population densities within two-species communities (Dunn’s test, *Z* = 4.48, *p*-adjusted < 0.05), and a pronounced 4.7-fold reduction in the three-species communities (Dunn’s test, *Z* = -8.62, *p*-adjusted < 0.05), relative to the near-isogenic control (Figure 2A). Subsequently, we considered the temporal changes as an influencing factor in population density and evaluated how population density changed between the two sampling points. Here we observed a small yet significant temporal effect on population density (Wilcoxon, *W* = 84,973, *p* < 0.05), with a 1.7-fold increase between 7 and 14 dpi.

**Figure 2.**
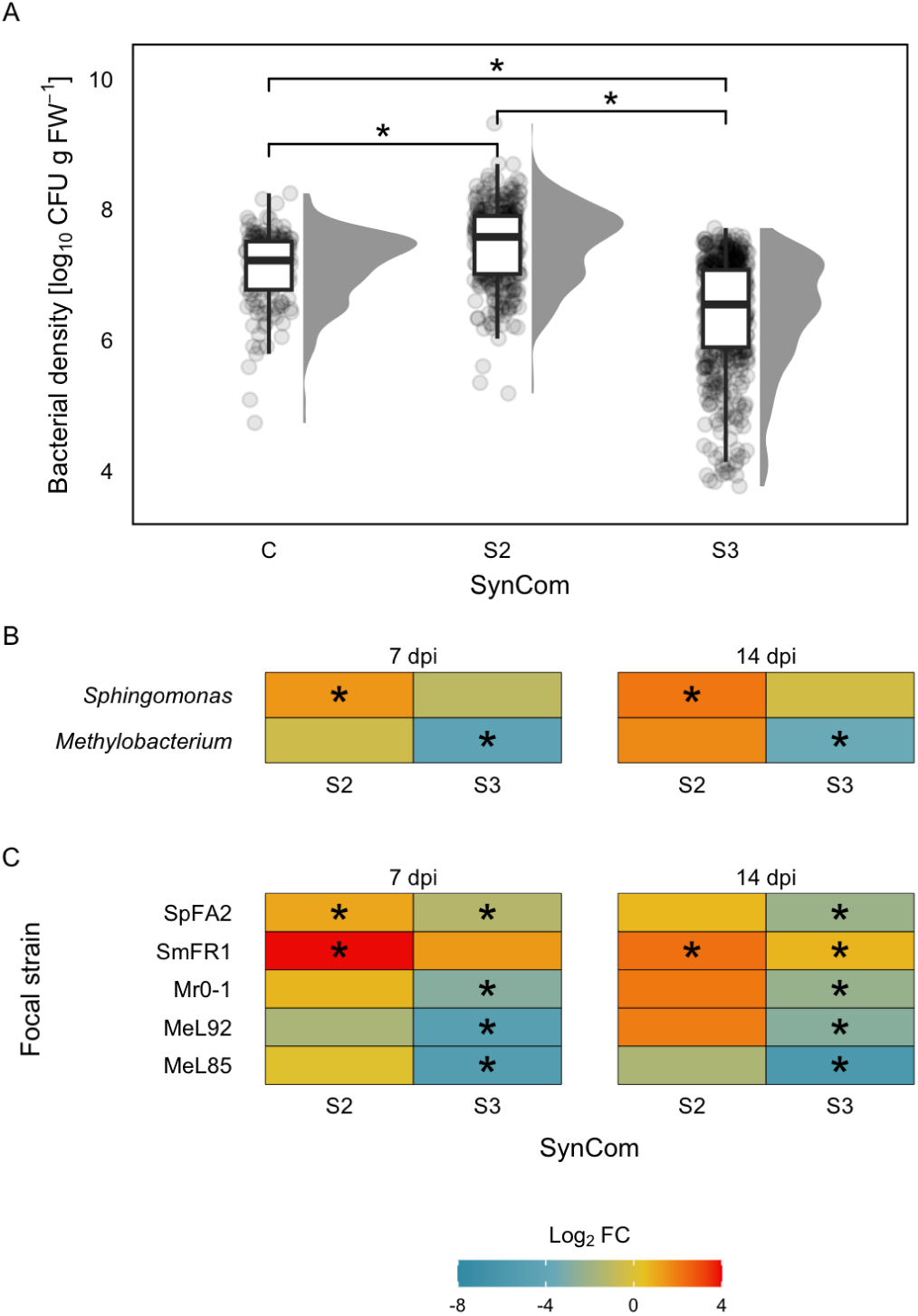
Bacterial population density in the arabidopsis phyllosphere. (A) Effect of community complexity (SynCom) on population densities. Asterisk (*) indicates statistical differences between groups based on Kruskal-Wallis and Dunn’s tests. *P*-values were corrected for multiple testing. (B) Fold change (Log_2_ FC) of population densities over time and in each SynCom per taxonomic group (*Methylobacterium* and *Sphingomonas*). (C) Fold change (Log_2_ FC) of population densities over time and in each treatment per strain. In B and C, fold change is relative to the near-isogenic control for each strain, and the asterisk indicates an adjusted *P*-value < 0.05.

Next, we evaluated the changes in populations of specific bacterial taxa, and found that *Methylobacterium* population densities were notably different to *Sphingomonas* (Wilcoxon, *Z* = 49,825.5, *p* < 0.05). *Sphingomonas* population densities were 5.1 times larger than those of *Methylobacterium* (Figure 2B). Within *Sphingomonas*, SmFR1 consistently increased population sizes in S2, irrespective of the presence of a second species, and to a lesser extent in S3 (Figure 2C, Figure S2). By contrast, SpFA2 population sizes showed a transient increase in S2, but predominantly decreased in S3 (Figure 2C).

*Methylobacterium* populations responded negatively to increasing community complexity (Figure 2B). MeL85, MeL92, and Mr0-1 consistently experienced a reduction in population sizes, particularly within S3 and, to a lesser extent, in S2 (Figure 2C). Among the *Methylobacterium* species, MeL92 and Mr0-1 benefited from the presence of any other species (S2). However, this effect was only observed at 14 dpi, and it was lost in the presence of a third competitor (S3). MeL85 was most susceptible to population decrease over time, and across varying community complexities and compositions (Figure S2).

Collectively, these findings indicate that bacterial taxa differentially responded to community complexity within the leaf environment. Notably, *Methylobacterium* populations were more susceptible compared to Sphingomonads. Among the studied species, *Sphingomonas* FR1 emerged as the most competitive, while *Methylobacterium* L85 was the least competitive.

### Spatial distribution of individual strains depends on their community context

We obtained an exemplary representation of every bacterial population at the single cell resolution, as well as their spatial distributions. This was accomplished by determining the centre of mass of individual cells within each bacterial combination (Figure S1A). The centres of mass of the different cell populations were then used to generate spatial point patterns and further used to analyse spatial distribution patterns of bacteria. In total, we analysed 10,261 fields of view, corresponding to 25 distinct communities (5× C, 10× S2, 10× S3), captured at two time points, with 3 to 4 biological replicates (individual plants). On average, each biological replicate was composed of 55 ± 30 fields of view (mean ± SD), in two independent experiments (Table S3).

We identified several characteristic patterns that follow the leaf surface topography, including veins and epidermal cell grooves, each with varying levels of occupancy (Figure 3). Subsequently, we observed that the cell distribution on the leaf surface was heterogenous by subsampling our data set to 500 random fields of view. We confirmed that these patterns can be described as a non-homogeneous Poisson process by a spatial Kolmogorov-Smirnov test (*D* = 0.04, *p* < 0.05), and a goodness-of-fit with Monte Carlo test (χ^2^ = 157,493, *p* < 0.05, n = 999).

**Figure 3.**
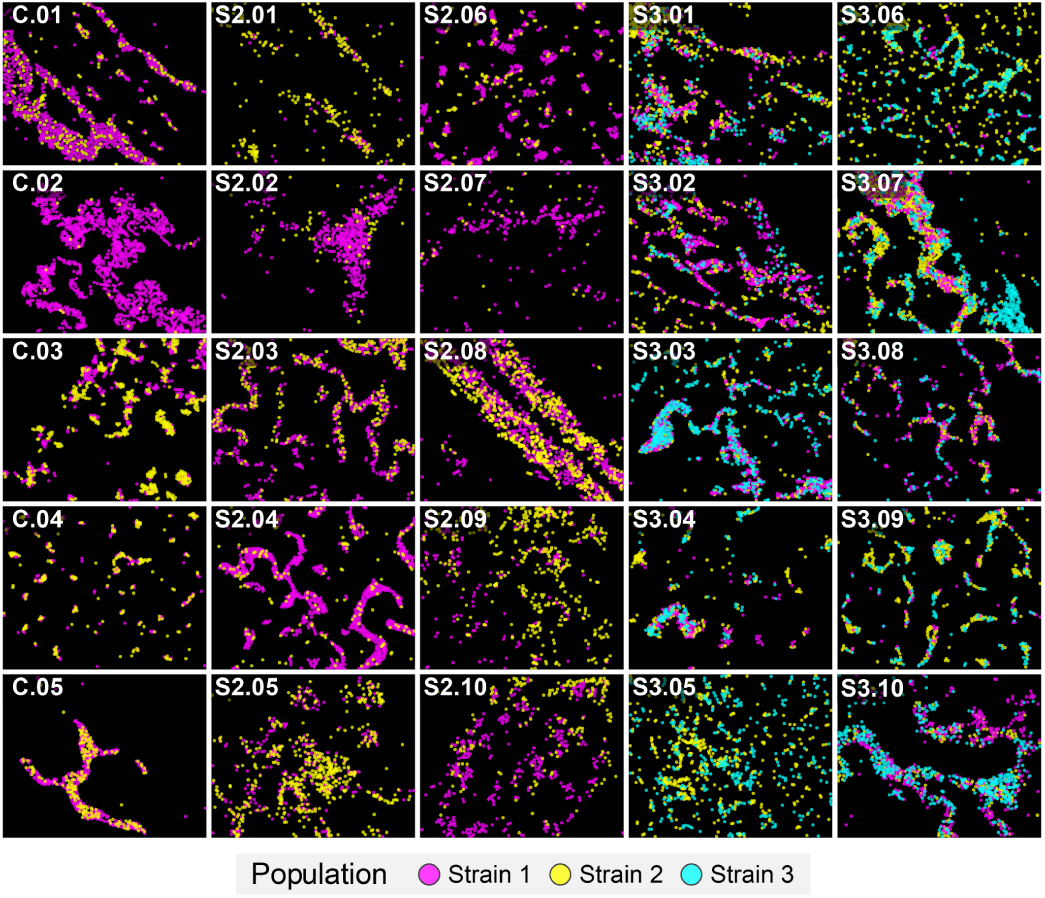
Representative images of point patterns from bacterial populations in the phyllosphere. Each image represents every tested bacterial combination (5 near-isogenic control C; 10 S2 communities; 10 S3 communities). Images were taken at 14 days post-inoculation. Refer to Table 2 for the identity of each strain in each combination.

Next, we estimated cell density per square centimetre for each biological replicate from single-cell microscopy data [# cell cm^-2^] and compared our estimations with colony count data [CFU gFW^-1^], which were positively correlated (Pearson’s *r* = 0.37, *t* = 11.55, *p* < 0.05). We observed a decrease in cell density between populations from S2 to S3 (Figure 4A). Differences in cell density were unrelated to time of sampling (Wilcoxon, *W* = 87,352, *p* = 0.56), but was rather driven by a decrease in *Methylobacterium* populations (log_2_FC = -1.08) and a marginal increase in *Sphingomonas* populations (log_2_FC = 0.34) (Figure 4B). These differences were associated with a decline in MeL85 populations and an increase in SmFR1 within S3 (Figure 4C). These observations indicate that the effect of the community complexity in a population was consistent at the CFU level and the single-cell resolution.

**Figure 4.**
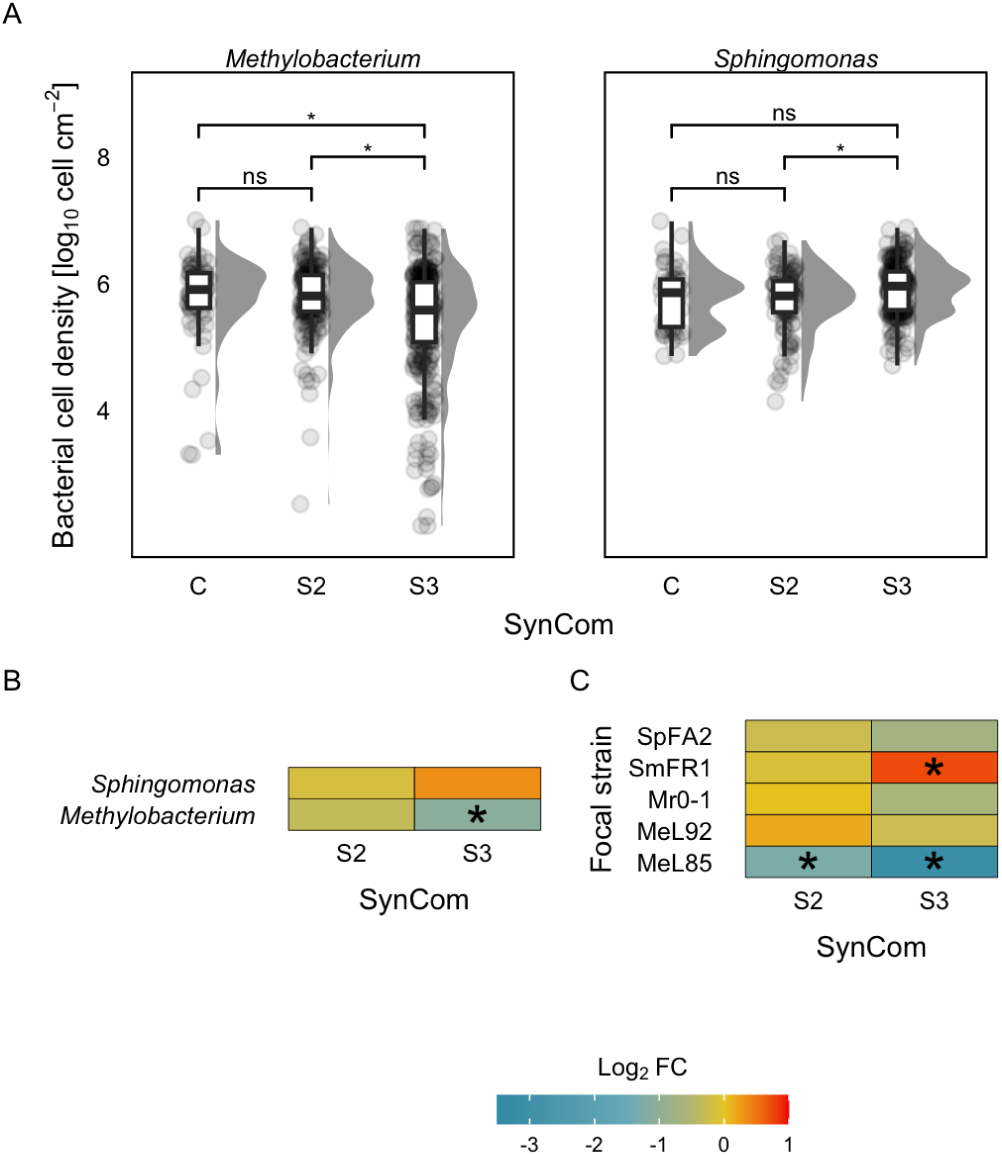
Bacterial cell densities in the phyllosphere. (A) Effect of community complexity (SynCom) on cell densities [log_10_ cell cm^-2^] for each taxonomic group (*Methylobacterium* and *Sphingomonas*). Asterisk (*) indicates statistical differences between groups based on Kruskal-Wallis followed by a post hoc Dunn’s test. *P*-values were corrected for multiple testing. (B) Fold change (Log_2_FC) of cell densities in each SynCom per taxonomic group. (C) Fold change (Log_2_FC) of cell densities in each treatment per strain. In B and C, fold change is relative to the near-isogenic control for each strain, and the circle size indicates the adjusted *P*-value from a one-sample Wilcoxon test (µ = 0, representing the near-isogenic control).

### Effect of community complexity on intraspecific spatial relations

Community context was expected to influence the spatial distribution patterns (aggregation, randomness, regularity) within bacterial populations in the phyllosphere. To evaluate this, we first determined relative frequencies of a spatial pattern based on ^K^^ (r), for each bacterial strain within every community context (Figure 5A, Figure S3). Subsequently, we quantified the area under the curve of each spatial pattern and calculated the fractional change compared to the near-isogenic control condition, C.

**Figure 5.**
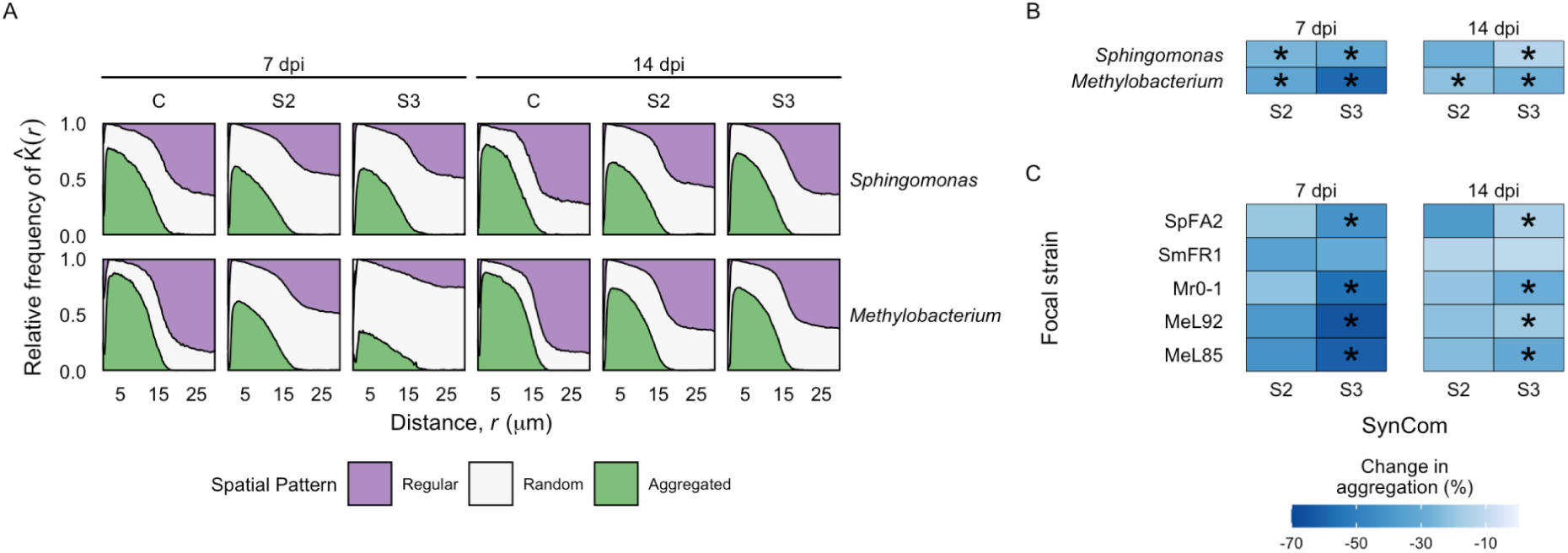
Intraspecific spatial patterns. (A) Frequency plots of spatial patterns determined by 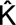(r) as a function of the distance between the centres of bacterial cells within a population. Regularity and aggregation suggests spatial segregation and clustering, respectively. Frequency plots of *Sphingomonas* and *Methylobacterium* groups within SynComs and time points. (B) Change in aggregation pattern (%) of a taxonomic group (*Sphingomonas* and *Methylobacterium*) relative to its near-isogenic control within S2 and S3 over time. (C) Change in aggregation pattern (%) of individual strains relative to its near-isogenic control within S2 and S3 over time. Circle size indicates the adjusted *P*-value from a one-sample Wilcoxon test (µ = 0, representing the near-isogenic control).

Our initial analysis showed that spatial distribution patterns within populations differed from their respective controls. Combined, there was a reduction of 27.5% and 26.1% in aggregation and regularity, respectively, while random distributions increased by 67.1% (Figure S4). Increase in randomness was observed mainly in *Methylobacterium*.

The observed distribution patterns were consistent across the two time points. That is, aggregation and regularity within populations decreased, while randomness increased in 7 and 14 dpi. Given that aggregation and regularity followed a consistent pattern, and randomness is a factor that would be related to neutral rather than deterministic processes, we decided to focus our analysis only on changes in aggregation within populations and the factors influencing this spatial pattern.

When comparing the aggregation between bacterial taxa, we identified differences between *Methylobacterium* and *Sphingomonas* only at 7 dpi (Wilcoxon, *W* = 133, *p* < 0.05). At this time point, aggregation differed from S2 to S3 (Wilcoxon, *W* = 452, *p* < 0.05). While both taxa exhibited decreased self-aggregation, *Methylobacterium* showed the largest decrease in aggregation in S3 (58.4%) relative to the control (Figure 5B). This decrease was reflected in a reduction of 61.5%, 65.1%, and 55.7% in aggregation for populations of MeL85, MeL92, and Mr0-1, respectively (Figure 5C). Conversely, the sphingomonads SmFR1 and SpFA2 were less affected, with reductions in self-aggregation of 29.4% and 40.9% in S3, respectively. Consequently, MeL85 and MeL92 were found to be statistically different to SmFR1 within S3.

The results of our analysis indicate that bacterial taxa respond differentially to community complexity, resulting in decreased self-aggregation and self-regularity, alongside an increase in random distribution patterns. This was particularly the case for *Methylobacterium* at 7 dpi in three-species communities. Furthermore and consistent with our results at the CFU and observed cell densities, aggregation was most affected in methylobacteria compared to sphingomonads, particularly MeL85 and MeL92 were negatively affected, while SmFR1 was least affected.

### Effect of community complexity on interspecific spatial correlations

We investigated the effect of community complexity on the spatial patterns between pairs of species. Pair cross correlations, ĝ(r), indicated whether an aggregated, random, or regular pattern was detected and its frequency for a species pair at a given distance (Figure 6A, Figure S5). Subsequently, we determined the area under the curve to compare spatial patterns between species pairs in S3 relative to S2. We treated S2 as our control, since S2 represents the pair of species without any other competitor present.

**Figure 6.**
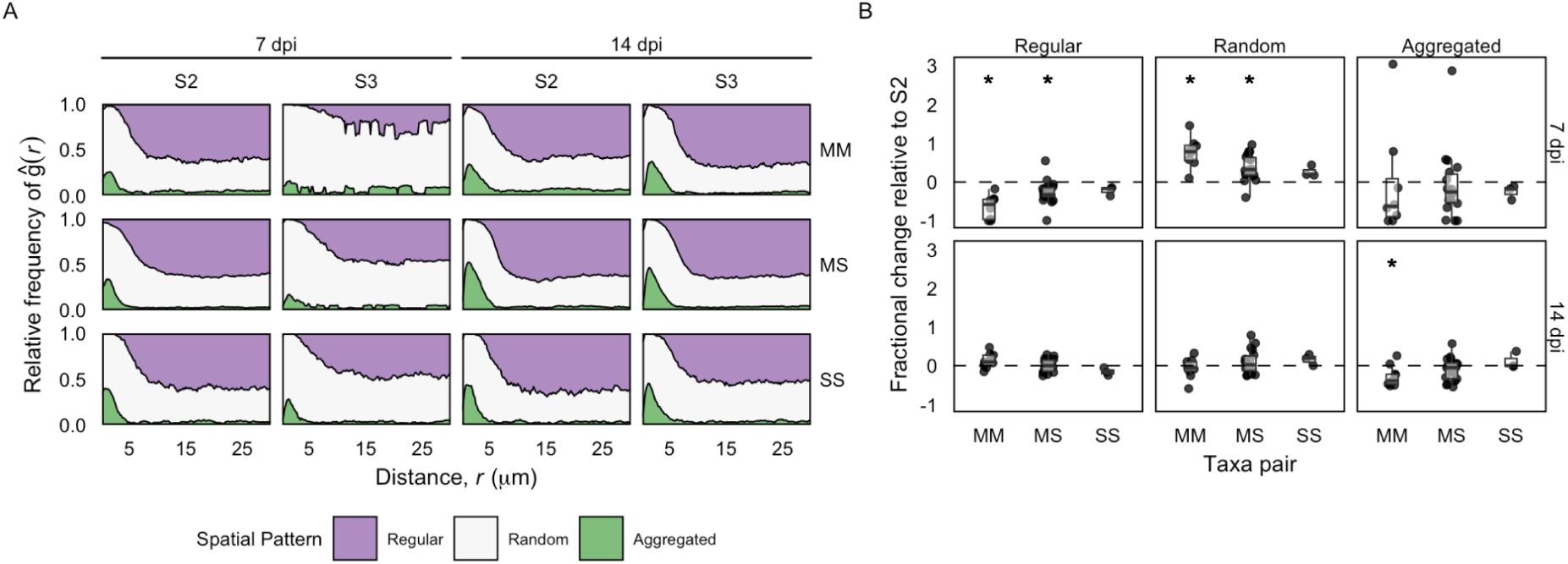
Interspecific spatial patterns. (A) Frequency plots of spatial patterns determined by pair cross correlation estimates as a function of the distance between the centres of bacterial cells from different populations. Combined plots for species pairs within *Methylobacterium* (MM), *Methylobacterium* and *Sphingomonas* (MS), and *Sphingomonas* (SS), grouped within SynComs and time points. (B) Fractional change of a spatial pattern (regular, random, aggregated) over time of a taxonomic pair (MM, MS, SS) relative to the same pair within S2. Asterisk (*) indicates statistical differences from a one-sample Wilcoxon test for a group of species pairs in S3 compared to the same pair in S2.

In general, we observed that pairs of species aggregated within S2 communities at each sampling point (Figure 6A). However, the presence of a third strain decreased these aggregation frequencies, subsequently altering regular and random patterns within S3 (Kruskal-Wallis, *H*(2) = 38.75, *p* < 0.05). Specifically, aggregation and regularity decreased by 18.9% and 15.4%, respectively, while randomness increased by 19.1%.

To further dissect these differences, we grouped the species pairs based on their identity, as a pair was composed of either two *Methylobacterium* (MM), a *Methylobacterium* and a *Sphingomonas* (MS), or two *Sphingomonas* (SS) species. We noticed that randomness was notably high at 7 dpi in pairs within S3 community (One-sample Wilcoxon, *W* = 422, *p* < 0.05), with this increase being particularly pronounced in pairs comprising at least one *Methylobacterium* species (MM and MS) (Figure 6B). However, the elevated randomness declined over time, becoming comparable to S2 at 14 dpi. The reduction in aggregation was primarily observed in MM pairs at 14 dpi. By contrast, pairs exclusively composed of *Sphingomonas* species (SS) were not different between S2 and S3.

Our findings collectively indicate that methylobacteria are more susceptible than sphingomonads to changes in their spatial distribution depending on their community context. This susceptibility is reflected by pairs containing at least one *Methylobacterium* species displaying random distribution patterns in more complex communities, implying that *Methylobacterium* tend to form fewer stable associations than *Sphingomonas*.

## Discussion

Microbial communities are shaped by neutral and niche factors, including stochastic processes, environmental conditions, and species interactions ^15,53^. In this study, we focused on the effects of community complexity as a factor influencing bacterial population densities and spatial distributions on the phylloplane of *Arabidopsis thaliana* for a group of specialist and generalist bacteria. Through a full factorial design involving near isogenic controls and SynComs of varying complexities (two-and three-species), we evaluated in detail the responses of *Methylobacterium* and *Sphingomonas* at different scales (bulk CFU and single cell). In general, our analysis showed significant effects of community composition and sampling time on bacterial population changes, with community complexity showing a higher influence than that of temporal dynamics.

Two-species communities showed a context-dependent increase in bacterial population densities compared to the near isogenic controls. This suggests that interactions between specific pairs of bacterial strains might lead to facilitation driven by mutualistic or parasitic relationships ^54^. The increase in population densities within these two-species communities was observed particularly in generalist sphingomonads, suggesting a fitness advantage of generalists over the specialist methylobacteria. Generalists can become dominant over specialists due to their metabolic flexibility and compensatory mechanisms ^55^. When community complexity was increased to three-species communities, bacterial population densities experienced a significant reduction, which was pronounced for methylobacteria. As *Methylobacterium* species mostly rely on methanol and only fewer other resources to sustain their growth ^21^, *Methylobacterium* populations are less versatiles in low diversity and local contexts, as observed for other specialists ^56^. Additionally, competition for a limiting factor such as methanol is expected to be strong among the three *Methylobacterium* species used in this study. We observed that MeL85 was the most susceptible strain, suggesting differential sensitivities of individual strains to the presence of competitors, even within a bacterial group. The observed variations in responses among *Methylobacterium* strains could arise from their distinct physiological characteristics, resource requirements, leaf colonisation capacity, and competitive abilities ^57–60^.

The contrasting responses of methylobacteria and sphingomonads to community complexity highlight the intricate nature of microbial interactions. Differences in two-species and three-species communities underscores the importance of considering higher-order interactions and the potential for shifting dynamics in more complex microbial communities ^61,62^. Stronger negative effects in population densities were observed in complex SynComs of three and seven members than those arising from pairwise interactions between leaf-associated bacteria ^21^. Higher-order interactions can reveal non-additive effects, as observed for SynComs with protective effects against a foliar pathogen ^18^ or, in contrast, SynComs associated with dysbiosis ^63^.

The spatial distribution of microbial populations plays a crucial role in understanding their ecological interactions and responses within complex communities. In this study, we examined the impact of community context on the spatial distribution of individual bacterial strains within the phyllosphere. By using single-cell resolution imaging and point pattern analysis, we revealed significant insights into how bacterial populations are arranged within simple communities. These spatial arrangements followed the leaf surface topography such as epidermal cell grooves, reflecting the influence of the physical microenvironment on bacterial positioning. Importantly, the observed distribution was not homogeneous, suggesting that bacterial colonisation does not occur randomly but adheres to an inhomogeneous Poisson process, common to heterogeneous environments ^13^. It is likely that instead of microbe-microbe interactions being a strong driver of spatial arrangements and community assembly, the topography and patchiness of the leaf surface have a stronger impact on spatial distributions of leaf colonisers. In this hypothetical, the spatial limitations force species to aggregate at a local scale. However, distinct spatial organisations can be driven by species interactions independent of the spatial context, as shown between *Pseudomonas fluorescens* or *Pseudomonas syringae* and the leaf-associated microbiota of their native hosts *ex situ*, i.e., flat surfaces ^64^. In our case, methylobacteria and sphingomonads exhibited distinct spatial patterns, indicating that the leaf topography alone cannot explain aggregation patterns between bacteria in this environment.

Furthermore, we validated our single-cell microscopy data by comparing cell density estimations with colony count data. This comparison revealed a positive correlation between the two methods, indicating the reliability of our single-cell imaging approach in estimating population densities. Differences in these estimations can be due to underestimation of total bacterial populations based on the leaf side ^65^, as we analysed populations from the abaxial side, and that not every cell might have been recovered from leaves ^12^. Despite discrepancies, similar trends in abundance changes between populations within complex communities were observed, highlighting consistent results from both bulk and single-cell resolutions.

In our analysis of the impact of community complexity on spatial relationships, we found that distribution patterns within and between populations changed in response to different community contexts. Notably, *Methylobacterium* strains exhibited reduced levels of intra-and interspecific aggregation within two-and three-species communities. This was consistent across different strains within *Methylobacterium*, with the most pronounced effect observed in the case of MeL85. Aggregation is a survival and stress response mechanism ^66–69^. Thus, increased randomness and decreased aggregation in *Methylobacterium* could be a reason for their population decline in more complex communities.

Conversely, *Sphingomonas* populations did not exhibit significant changes in aggregation patterns, indicating that responses to community complexity were taxon-specific. These changes in aggregation could be a result of strong adhesion and retention properties of bacteria to the leaf surface ^70^, facilitating clonal growth ^71^. At the same time, cell detachment from aggregates and motility are critical for population growth in the phyllosphere, as these processes enable new cells to explore uninhabited microenvironments ^9,72^. The significance of motility in conferring a fitness advantage in the phyllosphere is evident in *Pseudomonas* ^31,73^. However, this advantage should be further investigated for generalists and specialists. Notably, cell motility is a common trait among *Sphingomonas* ^74^, and is regulated during leaf colonisation, which is in contrast to the behaviour of *Methylobacterium* ^33^. Transcriptomic changes in genes related to motility have been reported in other leaf-associated bacteria ^75^, however, their relation to survival, growth, and interactions with other species are still unknown.

The ecological strategies of bacterial groups are linked to the balance between cell motility and aggregation in the phyllosphere. These strategies are manifested in the behaviour of *Sphingomonas* and *Methylobacterium*. *Sphingomonas* are fast-growing generalists, thus their metabolic flexibility could sustain larger mono and mixed aggregates from resource-depleted microenvironments. Concurrently, a fraction could migrate to richer sites, explaining the marginal reduction in self-aggregation, stable inter-species aggregation, and increased population size observed in this study.

By contrast, *Methylobacterium* are slow-growing specialist bacteria, and, although we expect migration to occur, their migration resulted in random distributions within complex communities (particularly S3). Coupled with their limited range of resource utilisation, a characteristic of specialists, and the relatively low diversity within S2 and S3 communities, which may not support species with limited metabolic flexibility, survival of *Methylobacterium* was compromised.

In summary, our study highlights the intricate responses of bacterial populations to community complexity within the phyllosphere and their spatial arrangements. The differential sensitivities of methylobacteria and sphingomonads and their distribution patterns we observed underscore the complexity of microbial interactions and their strategies to populate and coexist in the phyllosphere.

## Acknowledgements

We would like to thank Paula Jameson, Matthew Stott, and Matthias Rillig for insightful discussions. We would like to thank Evan Kear for experimental support. This work was supported by Marsden Fast Start grant number 17-UOC-057 (MRE). RS was supported by a New Zealand International Doctoral Research Scholarship (NZIDRS) and a University of Canterbury College of Science Ph.D. scholarship.

## Author Contributions

RS and MRE conceptualised the study. RS acquired and analysed the data. RS and MRE interpreted the data, and wrote the manuscript.

