## Supplemental Material for "Bacterial community complexity in the phyllosphere penalises specialists over generalists"

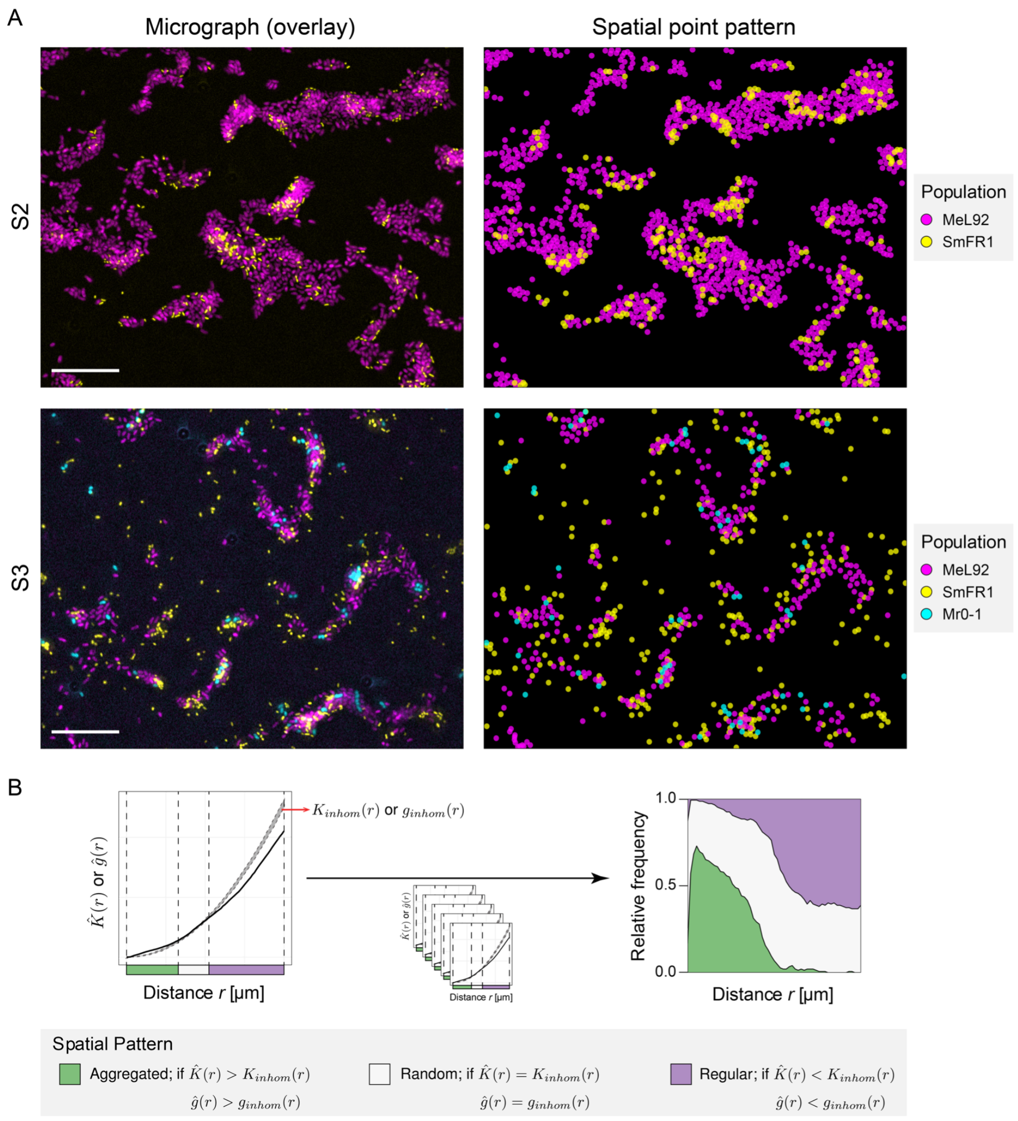


**Figure S1. Cell segmentation and spatial point pattern analysis.**

(A) *left:* Representative field of views of bacterial populations detected in an S2 (MeL92 and SmFR1) and S3 community (MeL92, SmFR1 and Mr0-1); *right:* corresponding spatial point pattern after cell segmentation and mapping of the cells in a micrograph. Thus, every point in the spatial point pattern represents the centre of mass of a bacterial cell. (B) For every point pattern, at a given distance r, either $\hat{K}(r)$ for intra-population or $\hat{g}(r)$ for inter-population spatial point patterns was estimated. A spatial aggregation pattern (green) was defined if $\hat{K}(r)$ or $\hat{g}(r)$ was greater than the upper limit of a Monte Carlo simulation envelope of the corresponding null models ($K_{inhom}(r)$ or $g_{inhom}(r)$, respectively). A random spatial pattern (grey) was defined if $\hat{K}(r)$ or $\hat{g}(r)$ fell within the Monte Carlo simulation envelope of the estimators. A regular or segregation pattern (purple) was defined if $\hat{K}(r)$ or $\hat{g}(r)$ was less than the lower limit of the Monte Carlo simulation envelope of the estimators. The frequency of each spatial pattern at a given distance was calculated based on the number of replicates and plotted for each tested condition.

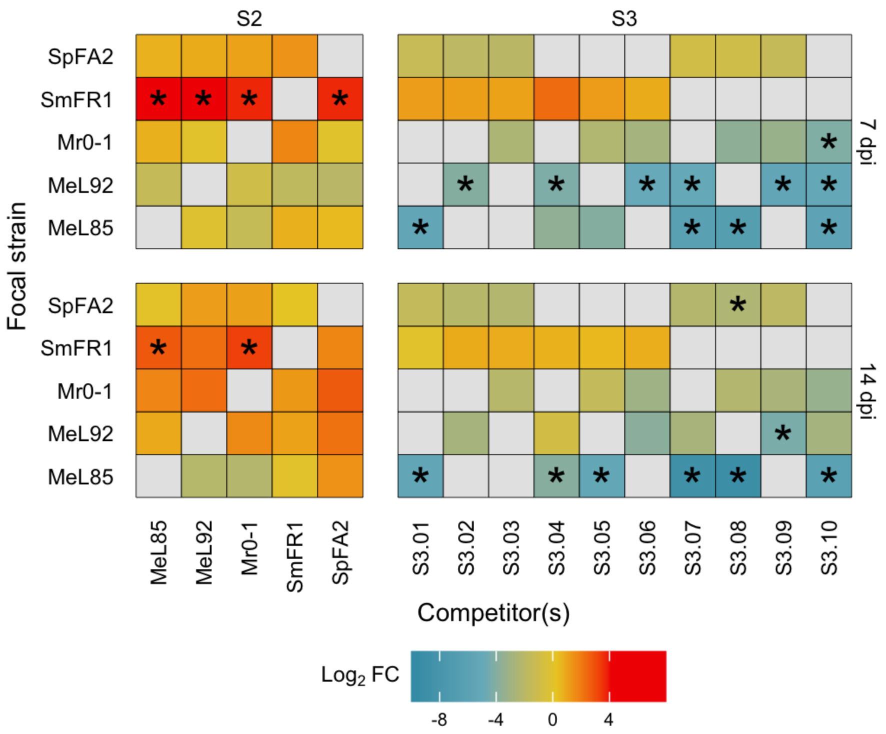


**Figure S2. Bacterial population densities per strains and in each community.** $Log_2$ fold change ($Log_2$ FC) of bacterial populations [CFU $gFW^{-1}$] in the presence of a single competitor (S2), or within S3. Fold changes were calculated from the median CFU of a strain in competition relative to the near-isogenic control. Asterisk indicates an adjusted *P*-value < 0.05, from a one-sample Wilcoxon test ($\mu = 0$, representing the near-isogenic control).

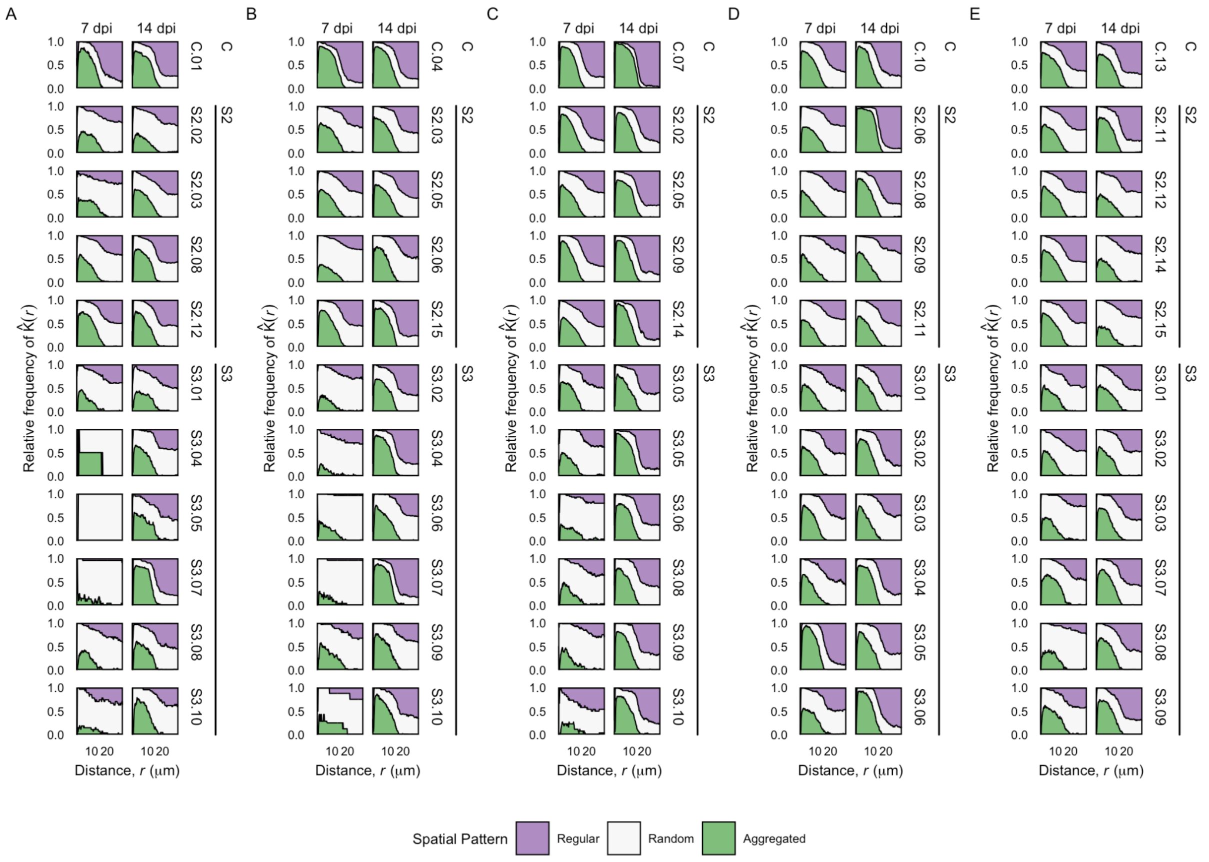


**Figure S3. Intraspecific frequency plots for each strain.**

Relative fractions of a spatial pattern (regular, random, aggregated) from Ripley's *K*-estimates for a bacterial population compared to the null model. (A) MeL85, (B) MeL92, (C) Mr0-1, (D) SmFR1, (E) SpFA2. Plots are grouped by community complexity (C, S2, S3) and time point. Refer to Table 2 in the main text for the identity of each strain in each combination.

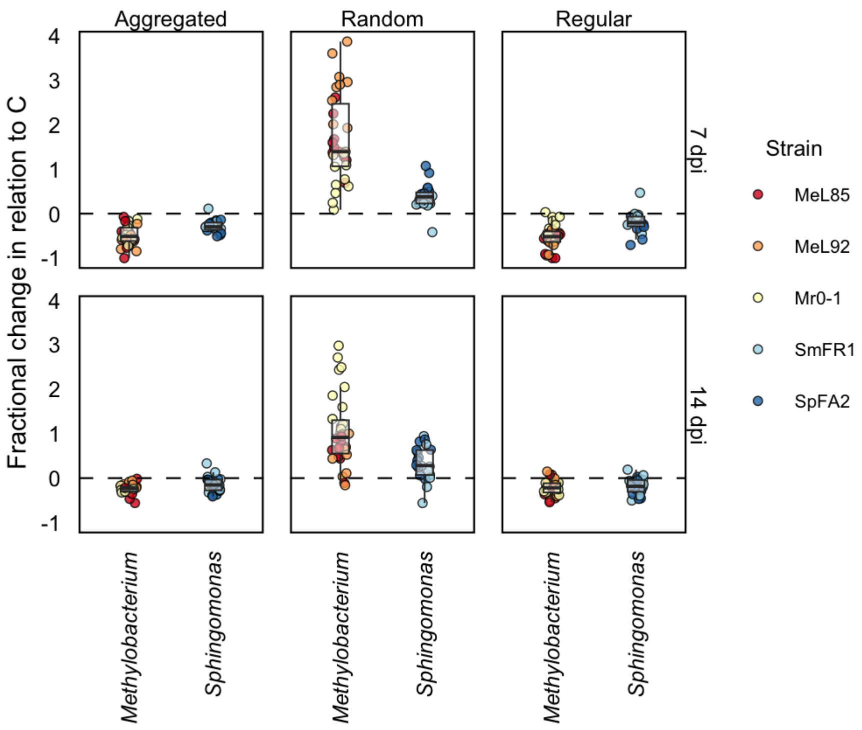


**Figure S4. Fractional change of a spatial pattern within bacterial taxonomic groups.**
Fractional change of a spatial pattern (aggregated, random, regular) over time of *Methylobacterium* and *Sphingomonas* relative to the corresponding near-isogenic control. Individual strains are highlighted within each group.

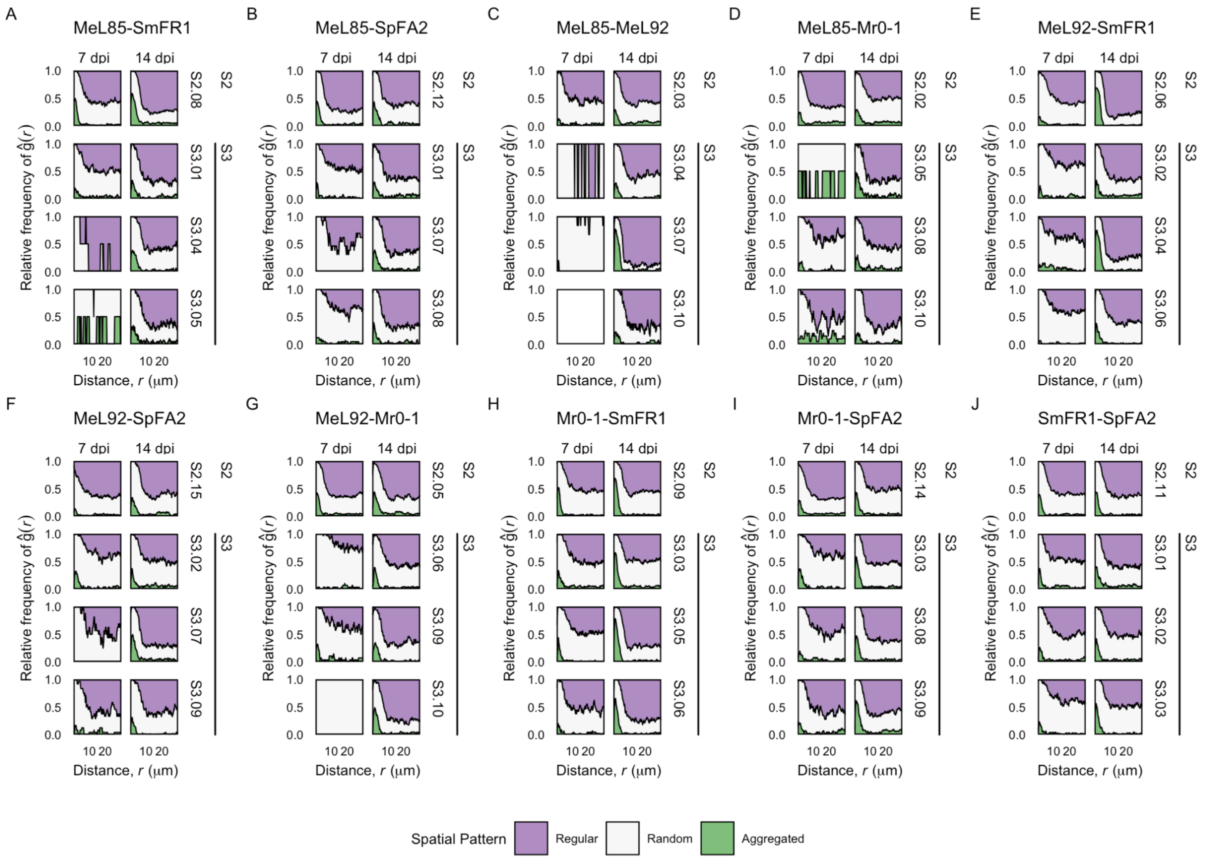


**Figure S5. Interspecific frequency plots for each species pair.**

Relative fractions of a spatial pattern (regular, random, aggregated) based on pair cross correlations between a bacterial pair compared to the null model. Frequency plots are grouped by bacterial pairs in S2 and S3, and time point. Species pair [Taxa pair]: (A) MeL85-SmFR1 [MS], (B) MeL85-SpFA2 [MS], (C) MeL85-MeL92 [MM], (D) MeL85-Mr0-1 [MM], (E) MeL92-SmFR1 [MS], (F) MeL92-SpFA2 [MS], (G) MeL92-Mr0-1 [MM], (H) Mr0-1-SmFR1 [MS], (**I**) Mr0-1-SpFA2 [MS], (J) SmFR1-SpFA2 [SS]. Refer to Table 2 in the main text for the identity of each strain in each combination.

**Table S1.** Strain selection within SynComs

| Treatment | Community | Strain | Fluorescence | Resistance | Selective media |
|---|---|---|---|---|---|
| C | C.01 | MeL85 | mSc | GmR | R2A + Gm |
| | | MeL85 | sYFP2 | KmR | R2A + Km |
| | C.02 | MeL92 | mSc | GmR | R2A + Gm |
| | | MeL92 | mTq2 | KmR | R2A + Km |
| | C.03 | Mr0-1 | mSc | GmR | R2A + Gm |
| | | Mr0-1 | mTq2 | KmR | R2A + Km |
| | C.04 | SmFR1 | mSc | GmR | NA + Gm |
| | | SmFR1 | mTq2 | TcR | NA + Tc |
| | C.05 | SpFA2 | mSc | GmR | NA + Gm |
| | | SpFA2 | mTq2 | TcR | NA + Tc |
| S2 | S2.01 | MeL85 | mSc | GmR | R2A + Gm |
| | | Mr0-1 | mTq2 | KmR | R2A + Km |
| | S2.02 | MeL85 | mSc | GmR | R2A + Gm |
| | | MeL92 | mTq2 | KmR | R2A + Km |
| | S2.03 | MeL92 | mSc | GmR | R2A + Gm |
| | | Mr0-1 | mTq2 | KmR | R2A + Km |
| | S2.04 | SmFR1 | mSc | GmR | NA + Gm |
| | | MeL92 | mTq2 | KmR | R2A + Km |
| | S2.05 | MeL85 | mSc | GmR | R2A + Gm |
| | | SmFR1 | mTq2 | TcR | NA + Tc |
| | S2.06 | Mr0-1 | mSc | GmR | R2A + Gm |
| | | SmFR1 | mTq2 | TcR | NA + Tc |
| | S2.07 | SpFA2 | mSc | GmR | NA + Gm |
| | | SmFR1 | mTq2 | TcR | NA + Tc |
| | S2.08 | MeL85 | mSc | GmR | R2A + Gm |
| | | SpFA2 | mTq2 | TcR | NA + Tc |
| | S2.09 | Mr0-1 | mSc | GmR | R2A + Gm |
| | | SpFA2 | mTq2 | TcR | NA + Tc |
| | S2.10 | MeL92 | mSc | GmR | R2A + Gm |
| | | SpFA2 | mTq2 | TcR | NA + Tc |
| S3 | S3.01 | SmFR1 | mSc | GmR | MM Proline + Gm |
| | | SpFA2 | sYFP2 | GmR | MM Serine + Gm |
| | | MeL85 | mTq2 | KmR | MM MeOH |
| | S3.02 | SmFR1 | mSc | GmR | MM Proline + Gm |
| | | SpFA2 | sYFP2 | GmR | MM Serine + Gm |
| | | MeL92 | mTq2 | KmR | MM MeOH |
| | S3.03 | SmFR1 | mSc | GmR | MM Proline + Gm |
| | | SpFA2 | sYFP2 | GmR | MM Serine + Gm |
| | | Mr0-1 | mTq2 | KmR | MM MeOH |
| | S3.04 | SmFR1 | mSc | GmR | MM Glucose + Gm |

| | | | | | |
|---|---|---|---|---|---|
| | | MeL85 | sYFP2 | KmR | MM Fructose + Km |
| | | MeL92 | mTq2 | KmR | MM MeOH + Km |
| | S3.05 | SmFR1 | mSc | GmR | MM Glucose + Gm |
| | | MeL85 | sYFP2 | KmR | MM Fructose + Km |
| | | Mr0-1 | mTq2 | KmR | MM MeOH + Km |
| | S3.06 | SmFR1 | mSc | GmR | MM Glucose + Gm |
| | | MeL92 | sYFP2 | GmR | MM MeOH + Gm |
| | | Mr0-1 | mTq2 | KmR | MM MeOH + Km |
| | S3.07 | MeL85 | mSc | GmR | MM MeOH + Gm |
| | | SpFA2 | sYFP2 | GmR | MM Glucose + Gm |
| | | MeL92 | mTq2 | KmR | MM MeOH + Km |
| | S3.08 | MeL85 | mSc | GmR | MM MeOH + Gm |
| | | SpFA2 | sYFP2 | GmR | MM Glucose + Gm |
| | | Mr0-1 | mTq2 | KmR | MM MeOH + Km |
| | S3.09 | MeL92 | mSc | GmR | MM MeOH + Gm |
| | | SpFA2 | sYFP2 | GmR | MM Glucose + Gm |
| | | Mr0-1 | mTq2 | KmR | MM MeOH + Km |
| | S3.10 | MeL85 | mSc | GmR | MM Fructose + Gm |
| | | MeL92 | sYFP2 | GmR | MM MeOH + Gm |
| | | Mr0-1 | mTq2 | KmR | MM MeOH + Km |

**Table S2.** Strain used in conjugation experiments

| Recipient strain | Donor strain | Phenotype(s) |
|---|---|---|
| *Sphingomonas melonis* Fr1 | *E. coli* S17-1 (pMRE-Tn5-153) | Yellow fluorescence (sYFP2)<br>Kanamycin resistance |
| *Sphingomonas phyllosphaerae* FA2 | *E. coli* S17-1 (pMRE-Tn5-143) | Yellow fluorescence (sYFP2)<br>Gentamicin resistance |
| *Sphingomonas phyllosphaerae* FA2 | *E. coli* S17-1 (pMRE-Tn5-161) | Cyan fluorescence (mTurquoise2)<br>Tetracycline resistance |
| *Methylobacterium radiotolerans* 0-1 | *E. coli* S17-1 (pMRE-Tn5-145) | Red fluorescence (mScarlet-I)<br>Gentamicin resistance |
| *Methylobacterium radiotolerans* 0-1 | *E. coli* S17-1 (pMRE-Tn5-153) | Yellow fluorescence (sYFP2)<br>Kanamycin resistance |
| *Methylobacterium radiotolerans* 0-1 | *E. coli* S17-1 (pMRE-Tn5-161) | Cyan fluorescence (mTurquoise2)<br>Tetracycline resistance |
| *Methylobacterium* sp. Leaf85 | *E. coli* S17-1 (pMRE-Tn5-145) | Red fluorescence (mScarlet-I)<br>Gentamicin resistance |
| *Methylobacterium* sp. Leaf85 | *E. coli* S17-1 (pMRE-Tn5-153) | Yellow fluorescence (sYFP2)<br>Kanamycin resistance |
| *Methylobacterium* sp. Leaf85 | *E. coli* S17-1 (pMRE-Tn5-151) | Cyan fluorescence (mTurquoise2)<br>Kanamycin resistance |
| *Methylobacterium* sp. Leaf92 | *E. coli* S17-1 (pMRE-Tn5-145) | Red fluorescence (mScarlet-I)<br>Gentamicin resistance |
| *Methylobacterium* sp. Leaf92 | *E. coli* S17-1 (pMRE105) | Yellow fluorescence (sYFP2)<br>Gentamicin resistance |
| *Methylobacterium* sp. Leaf92 | *E. coli* S17-1 (pMRE-Tn5-151) | Cyan fluorescence (mTurquoise2)<br>Kanamycin resistance |

**Table S3.** Summary of microscopy data.

| SynCom | Time of sampling [dpi] | Community | Indep. Exp. | nFOV | C0 [# cells] | C1 [# cells] | C2 [# cells] | Total [# cells] | Average [# cells/nFOV] |
|---|---|---|---|---|---|---|---|---|---|
| C | 7 | C.01 | e1 | 90 | 15719 | 64023 | n.a. | 79742 | 886.0 |
| | | C.01 | e2 | 10 | 128 | 1 | n.a. | 129 | 12.9 |
| | | C.02 | e1 | 203 | 24665 | 23976 | n.a. | 48641 | 239.6 |
| | | C.03 | e1 | 296 | 39755 | 25258 | n.a. | 65013 | 219.6 |
| | | C.04 | e1 | 192 | 18484 | 12200 | n.a. | 30684 | 159.8 |
| | | C.04 | e2 | 160 | 12739 | 23925 | n.a. | 36664 | 229.2 |
| | | C.05 | e1 | 210 | 11242 | 12336 | n.a. | 23578 | 112.3 |
| | | C.05 | e2 | 64 | 44230 | 17742 | n.a. | 61972 | 968.3 |
| | 14 | C.01 | e1 | 152 | 26928 | 9233 | n.a. | 36161 | 237.9 |
| | | C.01 | e2 | 80 | 23541 | 7415 | n.a. | 30956 | 387.0 |
| | | C.02 | e1 | 183 | 19205 | 20823 | n.a. | 40028 | 218.7 |
| | | C.02 | e2 | 81 | 13392 | 1095 | n.a. | 14487 | 178.9 |
| | | C.03 | e1 | 155 | 45063 | 32429 | n.a. | 77492 | 499.9 |
| | | C.04 | e1 | 104 | 17371 | 7771 | n.a. | 25142 | 241.8 |
| | | C.04 | e2 | 81 | 7642 | 10811 | n.a. | 18453 | 227.8 |
| | | C.05 | e1 | 121 | 16234 | 9300 | n.a. | 25534 | 211.0 |
| | | C.05 | e2 | 70 | 7517 | 9763 | n.a. | 17280 | 246.9 |
| S2 | 7 | S2.01 | e1 | 253 | 10175 | 45356 | n.a. | 55531 | 219.5 |
| | | S2.01 | e2 | 61 | 2836 | 11121 | n.a. | 13957 | 228.8 |
| | | S2.02 | e1 | 143 | 3705 | 10468 | n.a. | 14173 | 99.1 |
| | | S2.02 | e2 | 81 | 570 | 1873 | n.a. | 2443 | 30.2 |
| | | S2.03 | e1 | 207 | 24405 | 18589 | n.a. | 42994 | 207.7 |
| | | S2.03 | e2 | 130 | 6798 | 3411 | n.a. | 10209 | 78.5 |
| | | S2.04 | e1 | 203 | 6454 | 27234 | n.a. | 33688 | 166.0 |
| | | S2.04 | e2 | 163 | 9800 | 23432 | n.a. | 33232 | 203.9 |
| | | S2.05 | e1 | 226 | 3308 | 17081 | n.a. | 20389 | 90.2 |
| | | S2.05 | e2 | 81 | 14871 | 12478 | n.a. | 27349 | 337.6 |
| | | S2.06 | e1 | 250 | 36811 | 13552 | n.a. | 50363 | 201.5 |
| | | S2.06 | e2 | 81 | 20591 | 9408 | n.a. | 29999 | 370.4 |
| | | S2.07 | e1 | 251 | 14674 | 21637 | n.a. | 36311 | 144.7 |
| | | S2.07 | e2 | 80 | 14563 | 6963 | n.a. | 21526 | 269.1 |
| | | S2.08 | e1 | 187 | 3729 | 13363 | n.a. | 17092 | 91.4 |
| | | S2.08 | e2 | 80 | 13990 | 11083 | n.a. | 25073 | 313.4 |
| | | S2.09 | e1 | 300 | 28038 | 64385 | n.a. | 92423 | 308.1 |
| | | S2.09 | e2 | 164 | 16273 | 5759 | n.a. | 22032 | 134.3 |
| | | S2.10 | e1 | 220 | 26148 | 22799 | n.a. | 48947 | 222.5 |
| | | S2.10 | e2 | 163 | 22234 | 1934 | n.a. | 24168 | 148.3 |
| | 14 | S2.01 | e1 | 155 | 8458 | 45264 | n.a. | 53722 | 346.6 |
| | | S2.01 | e2 | 80 | 5947 | 15804 | n.a. | 21751 | 271.9 |
| | | S2.02 | e1 | 154 | 7890 | 27869 | n.a. | 35759 | 232.2 |
| | | S2.02 | e2 | 60 | 6679 | 2962 | n.a. | 9641 | 160.7 |
| | | S2.03 | e1 | 151 | 15246 | 20711 | n.a. | 35957 | 238.1 |

| | | | | | | | | | |
|---|---|---|---|---|---|---|---|---|---|
| | | S2.03 | e2 | 160 | 22661 | 71162 | n.a. | 93823 | 586.4 |
| | | S2.04 | e1 | 78 | 2011 | 10340 | n.a. | 12351 | 158.3 |
| | | S2.04 | e2 | 80 | 38332 | 11958 | n.a. | 50290 | 628.6 |
| | | S2.05 | e1 | 149 | 19727 | 25319 | n.a. | 45046 | 302.3 |
| | | S2.05 | e2 | 80 | 20104 | 21108 | n.a. | 41212 | 515.2 |
| | | S2.06 | e1 | 154 | 39837 | 15431 | n.a. | 55268 | 358.9 |
| | | S2.06 | e2 | 81 | 35256 | 15338 | n.a. | 50594 | 624.6 |
| | | S2.07 | e1 | 113 | 23671 | 20683 | n.a. | 44354 | 392.5 |
| | | S2.07 | e2 | 80 | 14687 | 6337 | n.a. | 21024 | 262.8 |
| | | S2.08 | e1 | 65 | 7479 | 12583 | n.a. | 20062 | 308.6 |
| | | S2.08 | e2 | 80 | 23042 | 10364 | n.a. | 33406 | 417.6 |
| | | S2.09 | e1 | 80 | 34940 | 8851 | n.a. | 43791 | 547.4 |
| | | S2.09 | e2 | 80 | 16125 | 12825 | n.a. | 28950 | 361.9 |
| | | S2.10 | e1 | 80 | 16312 | 4944 | n.a. | 21256 | 265.7 |
| | | S2.10 | e2 | 80 | 15255 | 4784 | n.a. | 20039 | 250.5 |
| S3 | 7 | S3.01 | e1 | 100 | 16524 | 6274 | 4650 | 27448 | 274.5 |
| | | S3.01 | e2 | 30 | 1634 | 2311 | 7 | 3952 | 131.7 |
| | | S3.02 | e1 | 80 | 5491 | 5907 | 2520 | 13918 | 174.0 |
| | | S3.02 | e2 | 80 | 11250 | 4081 | 2920 | 18251 | 228.1 |
| | | S3.03 | e1 | 80 | 13448 | 3868 | 17250 | 34566 | 432.1 |
| | | S3.03 | e2 | 77 | 13476 | 2054 | 2862 | 18392 | 238.9 |
| | | S3.04 | e1 | 81 | 14759 | 118 | 1305 | 16182 | 199.8 |
| | | S3.04 | e2 | 61 | 6758 | 8 | 636 | 7402 | 121.3 |
| | | S3.05 | e1 | 81 | 65369 | 81 | 13398 | 78848 | 973.4 |
| | | S3.05 | e2 | 60 | 16609 | 25 | 1693 | 18327 | 305.5 |
| | | S3.06 | e1 | 81 | 7453 | 1002 | 982 | 9437 | 116.5 |
| | | S3.06 | e2 | 60 | 10187 | 483 | 2876 | 13546 | 225.8 |
| | | S3.07 | e1 | 62 | 496 | 1643 | 395 | 2534 | 40.9 |
| | | S3.07 | e2 | 61 | 186 | 6830 | 276 | 7292 | 119.5 |
| | | S3.08 | e1 | 78 | 3473 | 2983 | 4094 | 10550 | 135.3 |
| | | S3.08 | e2 | 21 | 5 | 347 | 250 | 602 | 28.7 |
| | | S3.09 | e1 | 64 | 1253 | 1206 | 2008 | 4467 | 69.8 |
| | | S3.09 | e2 | 61 | 542 | 7422 | 1725 | 9689 | 158.8 |
| | | S3.10 | e1 | 41 | 1033 | 161 | 2164 | 3358 | 81.9 |
| | | S3.10 | e2 | 13 | 99 | 534 | 0 | 633 | 48.7 |
| | 14 | S3.01 | e1 | 81 | 10496 | 5378 | 12927 | 28801 | 355.6 |
| | | S3.01 | e2 | 60 | 10432 | 8820 | 2494 | 21746 | 362.4 |
| | | S3.02 | e1 | 80 | 23956 | 9518 | 19330 | 52804 | 660.1 |
| | | S3.02 | e2 | 60 | 14333 | 5609 | 3504 | 23446 | 390.8 |
| | | S3.03 | e1 | 81 | 10756 | 6064 | 13026 | 29846 | 368.5 |
| | | S3.03 | e2 | 60 | 8556 | 10008 | 34935 | 53499 | 891.7 |
| | | S3.04 | e1 | 80 | 13537 | 5659 | 10177 | 29373 | 367.2 |
| | | S3.04 | e2 | 60 | 11874 | 3298 | 14564 | 29736 | 495.6 |
| | | S3.05 | e1 | 80 | 13675 | 470 | 27362 | 41507 | 518.8 |
| | | S3.05 | e2 | 60 | 15034 | 2427 | 19273 | 36734 | 612.2 |
| | | S3.06 | e1 | 80 | 54483 | 4974 | 15092 | 74549 | 931.9 |
| | | S3.06 | e2 | 60 | 11573 | 9679 | 11861 | 33113 | 551.9 |
| | | S3.07 | e1 | 80 | 20166 | 8322 | 22215 | 50703 | 633.8 |

| | | | | | | | | | |
|---|---|---|---|---|---|---|---|---|---|
| | | S3.07 | e2 | 60 | 507 | 14232 | 13617 | 28356 | 472.6 |
| | | S3.08 | e1 | 80 | 2718 | 8688 | 23567 | 34973 | 437.2 |
| | | S3.08 | e2 | 61 | 4809 | 7009 | 3567 | 15385 | 252.2 |
| | | S3.09 | e1 | 81 | 5032 | 9833 | 25215 | 40080 | 494.8 |
| | | S3.09 | e2 | 60 | 1010 | 9160 | 8604 | 18774 | 312.9 |
| | | S3.10 | e1 | 50 | 2722 | 3286 | 8123 | 14131 | 282.6 |
| | | S3.10 | e2 | 59 | 26 | 15367 | 12899 | 28292 | 479.5 |

e1: Independent experiment #1
e2: Independent experiment #1
nFOV: Number of field of views
